# CIB2 function is distinct from Whirlin in the development of cochlear stereocilia staircase pattern

**DOI:** 10.1101/2024.07.30.605852

**Authors:** Arnaud P. J. Giese, Andrew Parker, Sakina Rehman, Steve D. M. Brown, Saima Riazuddin, Craig W. Vander Kooi, Michael R. Bowl, Zubair M. Ahmed

## Abstract

Variations in genes coding for calcium and integrin binding protein 2 (CIB2) and whirlin cause deafness both in humans and mice. We previously reported that CIB2 binds to whirlin, and is essential for normal staircase architecture of auditory hair cells stereocilia. Here, we refine the interacting domains between these proteins and provide evidence that both proteins have distinct role in the development and organization of stereocilia bundles required for auditory transduction. Using a series of CIB2 and whirlin deletion constructs and nanoscale pulldown (NanoSPD) assays, we localized the regions of CIB2 that are critical for interaction with whirlin. AlphaFold 2 multimer, independently identified the same interacting regions between CIB2 and whirlin proteins, providing a detailed structural model of the interaction between the CIB2 EF2 domain and whirlin HHD2 domain. Next, we investigated genetic interaction between murine *Cib2* and *Whrn* using genetic approaches. Hearing in mice double heterozygous for functionally null alleles (*Cib2^KO/+^;Whrn^wi/+^*) was similar to age-matched wild type mice, indicating that partial deficiency for both *Cib2* and *Whrn* does not impair hearing. Double homozygous mutant mice (*Cib2^KO/KO^;Whrn^wi/wi^*) had profound hearing loss and cochlear stereocilia exhibited a predominant phenotype seen in single *Whrn^wi/wi^* mutants. Furthermore, over-expression of *Whrn* in *Cib2^KO/KO^*mice did not rescue the stereocilia morphology. These data suggest that, CIB2 is multifunctional, with key independent functions in development and/or maintenance of stereocilia staircase pattern in auditory hair cells.

## Introduction

Hearing depends upon hair cells, the polarized epithelial cells of the inner ear that have mechano-sensitive hair bundles located at their apical pole. The hair bundle is composed of numerous stereocilia that are organized in a graded staircase pattern. The staircase architecture of the stereocilia bundle is conserved across all vertebrate hair cells and essential for hearing function, because it allows effective pulling of the tip links between stereocilia of neighboring rows. The organization, elongation and row identity of stereocilia in cochlear hair cells are highly regulated and involve several protein complexes, including MYO3A, MYO3B^1^, MYOXVA^2^, whirlin^3^, Eps8^4,5^, Eps8- L2^6^, and Gpsm2/Gα i3^7–9^. For instance, a long isoform of whirlin, is localized at the very tips of stereocilia and, together with its carrier – Myosin-XVa, is essential for the normal elongation of stereocilia and formation of the characteristic staircase shape of the hair bundle^2,3^. Furthermore, during development, whirlin targets Gpsm2/Gα i3 to the tips of first row stereocilia leading to the accumulation of the elongation protein complex at this site. In the absence of Gpsm2/Gα i3 complex, the first row stereocilia fail to grow to the correct height, resulting in profound deafness^7–9^.

We previously reported that in calcium and integrin binding protein 2, encoded by *Cib2*, homozygous mutant mice, the overall architecture of the cochlear stereociliary bundle is affected. CIB2 deficiency results in overgrowth of transducing shorter row stereocilia (rows 2 and 3) in the auditory hair cells without affecting non-transducing tallest row stereocilia (row 1), suggesting a direct role of CIB2 in stereocilia staircase patterning^10^. We have also demon- strated that CIB2 physically interacts with whirlin^11^. Given the role of both CIB2 and whirlin in the normal staircase patterning of stereocilia, and their binding with each other, we sought to determine a) their interacting domains; b) if CIB2 and whirlin have functional overlap with each other; and c) if there is genetic interaction between the genes encoding for CIB2 and whirlin. To map the interacting regions, we generated a series of CIB2 and whirlin deletion and point mutations fluorescently tagged constructs, performed nanoscale pull down interaction assays (Nano-SPD), co-immunoprecipitation studies, and molecular modeling. For functional interactions, we adapted classical genetic approaches, and crossed *Cib2^KO^* mice with *Whrn*^wi^ (knockout) or *Whrn^BAC279^* (over-expresser)^12^ mice and analyzed their first- and second-generation offspring. Analysis of double heterozygotes and double homozygous mutants allowed us to determine whether loss of CIB2 and whirlin affects viability, as both proteins are expressed in many tissues besides inner ear^11,12^, and produces a more severe inner ear phenotype or a superimposition of pathologies. Analysis of double heterozygotes also allowed us to determine digenic interaction between *Cib2* and *Whrn*. Finally, analysis of offspring from *Cib2^KO/KO^*, *Whrn^BAC279^*crosses was used to determine if over-expressing whirlin could rescue the stereocilia staircase pathology. Our data suggest that CIB2 has a role that is distinct from whirlin in development and/or organization of the stereocilia staircase patterning in the auditory hair cells.

## Results

### CIB2 is essential for the auditory hair cell stereocilia staircase pattern

Our previous studies revealed that the row identity of the cochlear stereociliary bundles was not maintained in *Cib2* mutant mice^10^. Here, we investigated if the observed loss of row identity is due to a role of CIB2 in the regulation of proteins essential for the staircase pattern (e.g. MyoXVA, whirlin, EPS8 etc.)^2–5^. To test this hypothesis, we immunostained organs of Corti from *Cib2* mutant mice (*Cib2^KO/KO^*) along with controls at P12, for the proteins, MyoXVa, whirlin, EPS8, and EPS8L2. Immunolabelling using PB48 antibody revealed aberrant staining pattern for Myosin XVa in *Cib2^KO/KO^* mutants, with over-accumulation at the tips of first and second row stereocilia of inner hair cell (IHC) bundles (Figure 1A). The overaccumulation of Myosin XVa was further confirmed by the quantification of fluorescent signal measured using confocal microscopy (Figure 1B). Furthermore, in the IHCs of *Cib2^KO/KO^* mice, whirlin immunostaining, detected using antibodies specific to long isoform of whirlin^3^, was weaker at the tips of stereocilia (Figure 1A-B). However, EPS8 and EPS8L2 immunostaining in the IHCs of *Cib2^KO/KO^* mice persisted at levels similar to that observed at the tips of stereocilia of control hair cells (Figure 1A-B).

**Figure 1:**
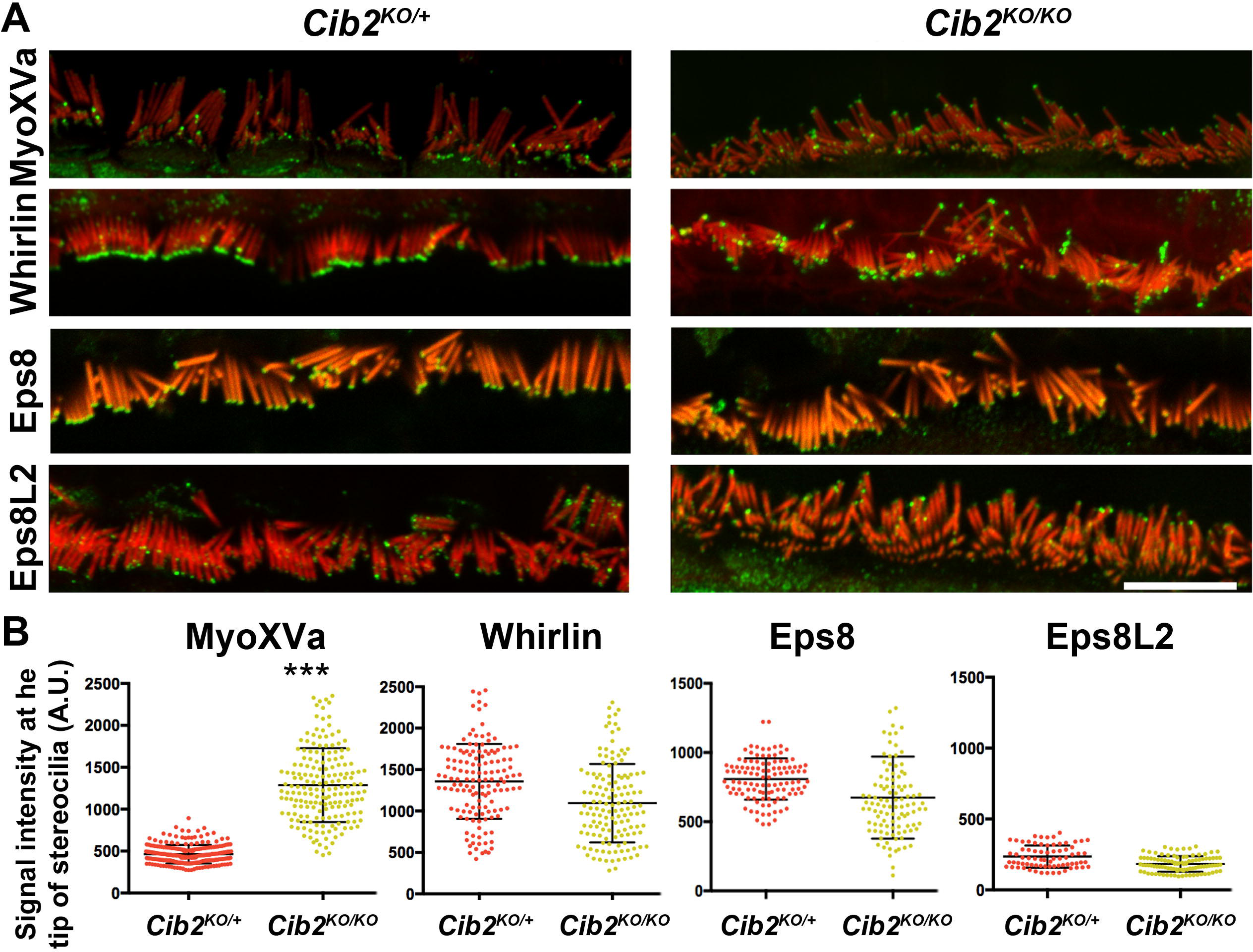
MyoXVa over accumulates at the tips of stereocilia in *Cib2^KO/KO^*mice. **A.** Expression of MyoXVa, whirlin, EPS8, and EPS8L2 proteins in *Cib2^KO/KO^* organs of Corti along with controls at P12. In contrast to other proteins, Myosin XVa was over-accumulation at the tip of the first and second row of stereociliary bundle of IHCs. **B.** Quantification of fluorescent signal measured by confocal microscopy at the tips of stereocilia, which further confirmed overaccumulation of Myosin XVa (\*\*\**p*<0.005).

### CIB2 EF2 binding motif is necessary for CIB2-whirlin interaction

We previously documented that CIB2 directly interacts with whirlin, and forms a tri-partite complex with whirlin and MyoXVa^11^. To further confirm these findings and to characterize the specific domains required for the CIB2- whirlin interaction, we performed nanoscale pulldown assays (NanoSPD)^13^.

For these studies, COS-7 cells were co-transfected with GFP- whirlin and various mCherry-Myo10-CIB2 deletion constructs (Figure 2A-B; Figure S1). The assay, using mCherry-Myo10-CIB2 and full-length GFP-whirlin constructs, confirmed interaction of these proteins. Further, both proteins significantly accumulated at the tip of filopodia in COS-7 cells as compared to negative control cells transfected with either mCherry-Myo10 or GFP-whirlin only (Figure 2B-C).

**Figure 2:**
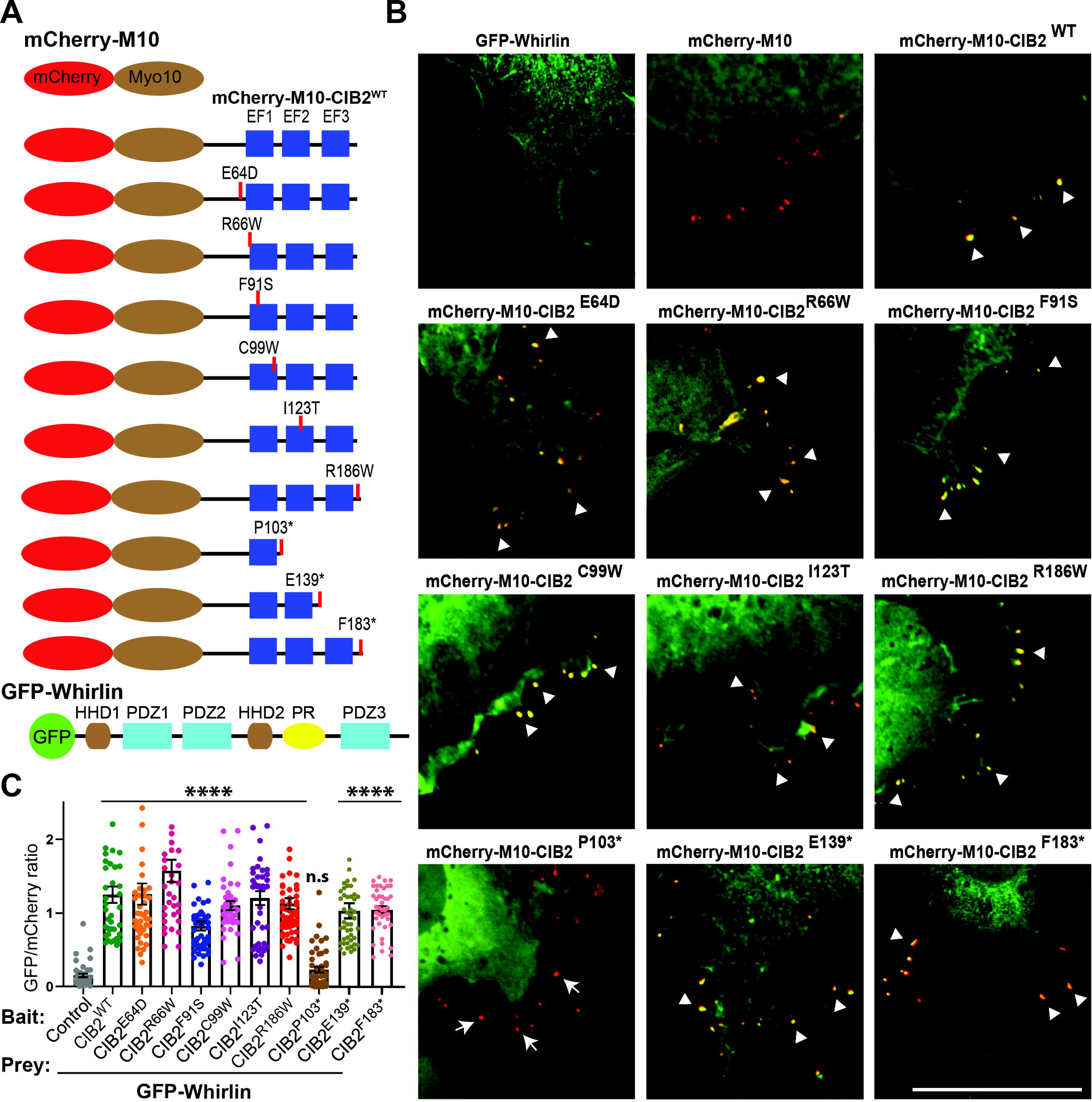
CIB2 EF2 domain binds Whirlin. **A.** Schematic of the mCherry-myo10, mCherry-myo10-CIB2^WT^ and CIB2 variants harboring, as well as GFP-whirlin^WT^ constructs used for the nanoSPD 1.0 assay. **B.** COS-7 cells were co-transfected with mCherry-myo10 or mCherry-myo10-CIB2 constructs (Baits, red) and GFP-whirlin (Prey, green), Merge channels are shown, and please see supplementary Figure S1 for single channel images. Accumulations at the tip of bait and prey are shown with an arrowhead. Arrows indicate the absence of accumulation of prey at the filopodia tip. Scale bar = 10 μm. **C.** Quantification of Nanoscale pulldown assay showing the interaction between whirlin and different CIB2 mutated constructs carrying some pathogenic DFNB48 missense variants, as well as truncations. \*\*\*\**p*≤ 0.0001; n.s = non-significant.

Next, we performed NanoSPD assays in COS-7 cells using various mCherry-Myo10-CIB2 deafness-causing missense variants (p.Glu64Asp, p.Arg66Trp, p.Phe91Ser, p.Cys99Trp, p.Ile123Thr, p.Arg186Trp) and GFP- whirlin full-length constructs (Figure 3B-C, Figure S1). As shown in Figure 3C, none of the missense pathogenic variants affect the CIB2-whirlin complex. As bait, we then used several truncated CIB2 constructs deleting the EF hand domains (p.Pro103*, p.Gln139*, p.Phe183*). The p.Pro103* truncation completely abolished the whirlin interaction, while p.Gln139* and p.Phe183* truncated CIB2 proteins retain the affinity of whirlin (Figure 3B-C).

**Figure 3:**
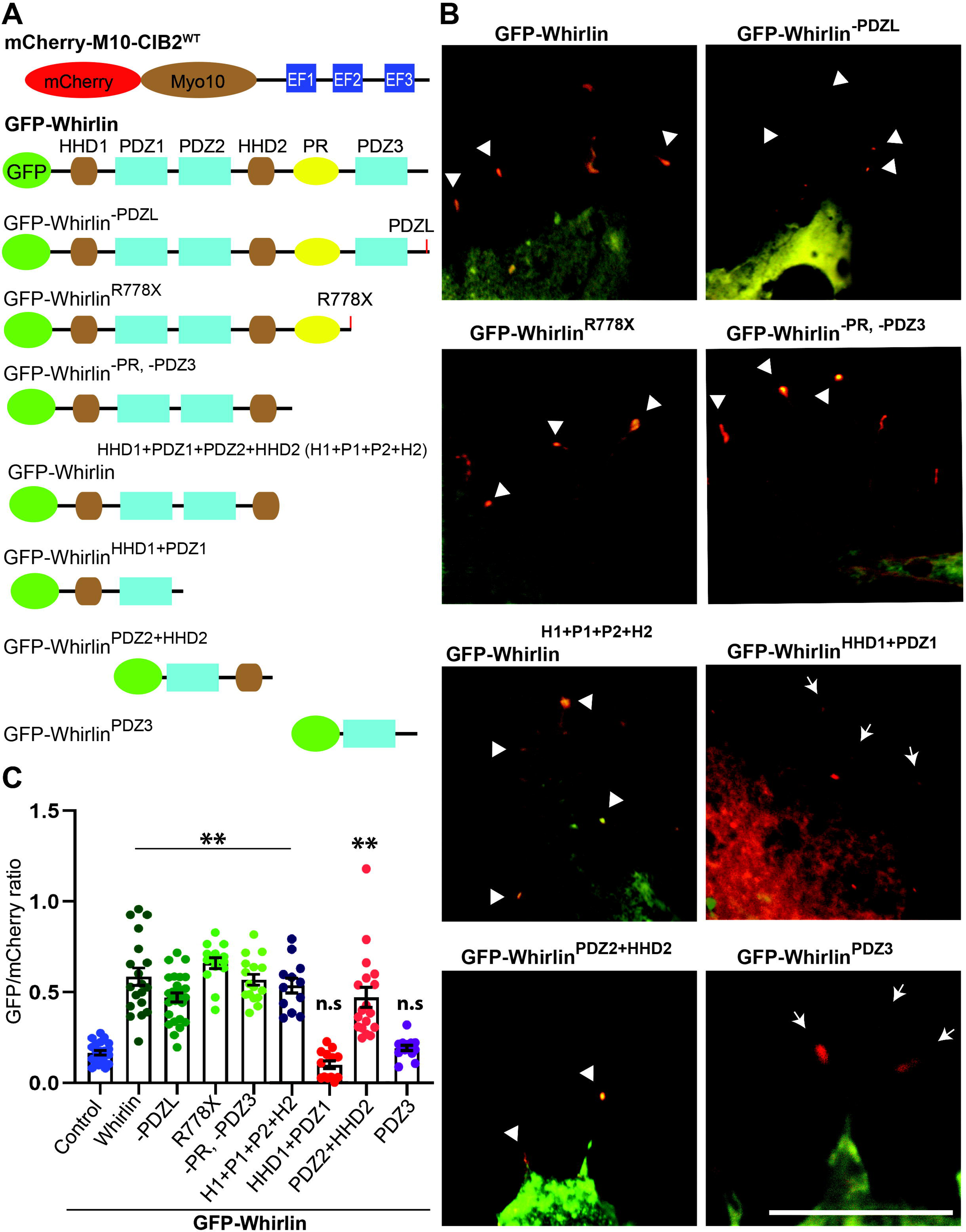
Whirlin PDZ2-HDD2 regions bind CIB2. **A.** Schematic of the mCherry-myo10-CIB2^WT^ and GFP-whirlin^WT^ and deletion constructs used for the nanoSPD 1.0 assay. **B.** COS-7 cells were co-transfected with mCherry-myo10-CIB2^WT^ (Bait, red) and GFP whirlin constructs (Preys, green), Merge channels are shown, and please see supplementary Figure S2 for single channel images. Accumulations at the tip of bait and prey are shown with an arrowhead. Arrows indicate the absence of accumulation of prey at the filopodia tip. Scale bar = 10 μm. **C.** Quantification of Nanoscale pulldown assay showing the interaction between whirlin PDZ2-HDD2 region and CIB2. \*\*\*\**p*≤ 0.0001; n.s = non-significant.

### Whirlin HDD2 domain is required for interaction with CIB2

We next investigated the specific domain region within whirlin required for interaction with CIB2. We performed NanoSPD assay in COS-7 cells using full-length mCherry-Myo10-CIB2 and GFP-whirlin truncated constructs (Figure 3, and Figure S2). All the GFP-whirlin truncated constructs that have PDZ2 and HDD2 domains in them co-accumulated at the tip of filopodia suggesting that the PDZ1 and PDZ3 domains may not be necessary for CIB2-whirlin interaction (Figure 3B-C, S2).

To gain detailed insight into the interacting regions of CIB2-whirlin, we generated an AlphaFold 2-multimer (AF2)^14,15^ model of the CIB2 in complex with whirlin (Figure 4A, and Figure S3). Significant complex formation was predicted between CIB2 and well-defined regions of whirlin. The predicted interaction is between the CIB2 C-terminal EF2 domain and the HDD2 of whirlin (Figure 4A). The involvement of the CIB2 C-terminal domain is consistent with deletion experiments, discussed above. Further, analysis of the complex indicated that it was strongly centered on the HDD2 domain of whirlin, with eight of ten potential salt bridges and twelve of sixteen potential hydrogen bonds contributed by the HDD2 domain of whirlin (Figure S3).

**Figure 4:**
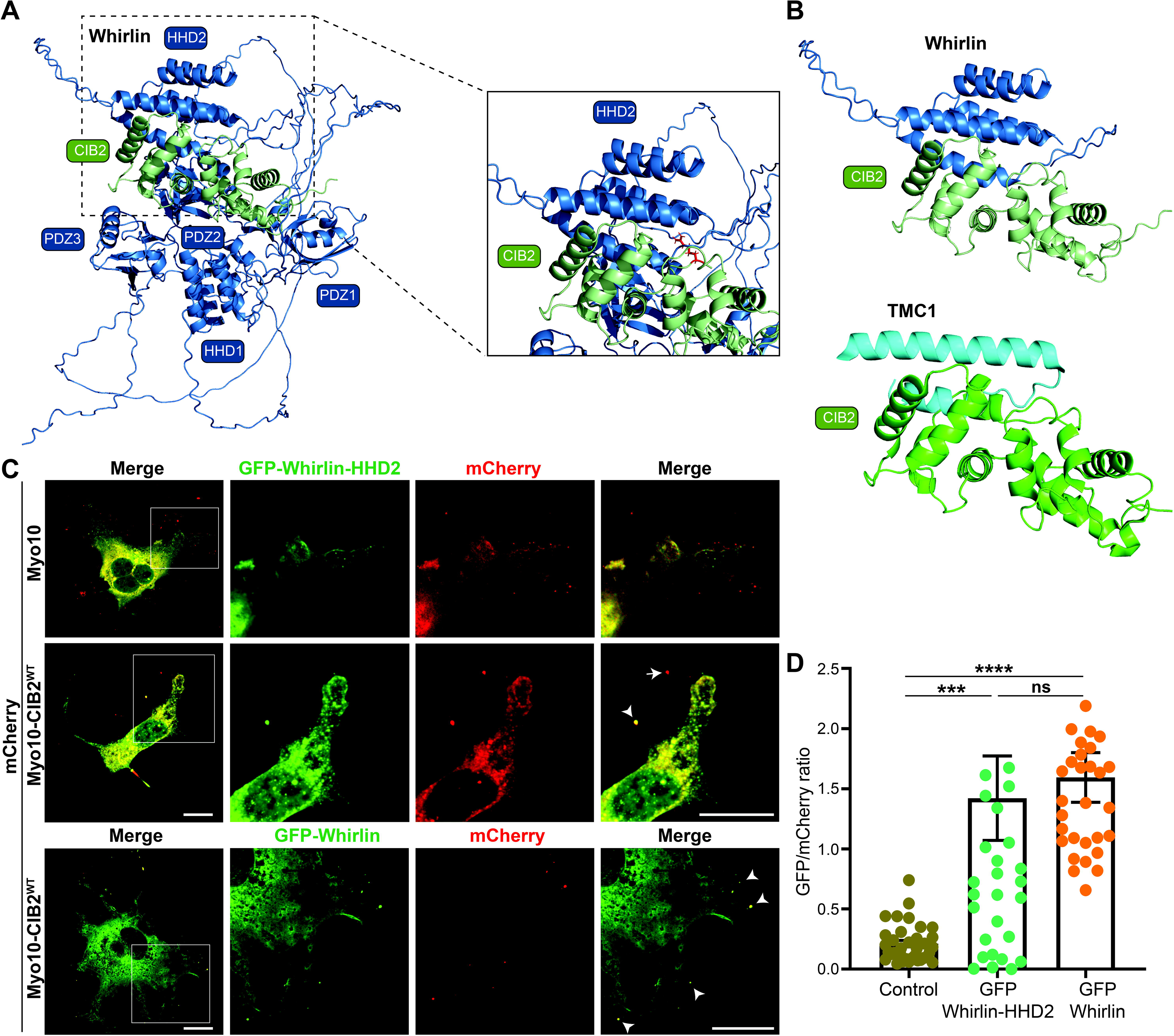
Whirlin HDD2 region binds CIB2 EF2 region. **A.** Alphafold multimer model of the complex between whirlin (blue) and CIB2 (green). **B.** Zoom of the predicted specific interacting HHD2 region of CIB2 (upper) and comparison to the known structure of TMC1 bound to CIB2 (lower). **C**. COS-7 cells were co-transfected with mCherry-myo10-CIB2^WT^ (Bait, red) and GFP-whirlin-HDD2 construct (Prey, green). Merge channels for whole cell, while single and merged channels for zoom in (boxed) regions are shown. Accumulations at the tip of bait and prey are shown with an arrowhead. Scale bar = 10 μm. **D.** Quantification of Nanoscale pulldown assay showing the interaction between whirlin PDZ2-HDD2 region and CIB2. \*\*\**p*≤ 0.001; \*\*\*\**p*≤ 0.0001, n.s = non-significant.

Strikingly, this potential interaction is highly structurally homologous to that formed between the high affinity TMC1 binding-region and CIB2^16^ (Figure 4B). Indeed, the core helical CIB2 binding regions have shared highly hydrophobic faces bound to the EF2 domain of CIB2.

Based on these predictions, we generated a GFP-whirlin-HDD2 domain only construct (Acc# NM_001008791.2; residues 415-561) and tested its interaction with full-length CIB2 through NanoSPD assay (Figure 4C).

Consistent with the Alphafold prediction, CIB2 interaction with HDD2 region was comparable to full-length whirlin protein (Figure 4D). Taken together, these data establish that the CIB2-whirlin interaction is mediated through the EF2 domain of CIB2 and the HDD2 domain of whirlin.

### Altering whirlin levels does not restore normal stereocilia architecture in *Cib2^KO/KO^* mice

Given the interaction between CIB2-whirlin, increased expression of MyoXVa (whirlin transporter) in *Cib2* mutant mice, and role of both proteins in orchestrating the staircase pattern of stereocilia bundle, next we sought to determine if altered whirlin levels are responsible for impaired stereocilia architecture in *Cib2^KO/KO^* mice using a classical genetic approach. To test this, we crossed *Cib2^KO/KO^* with *Whrn^wi/wi^* (Figure 5A) and analyzed first- and second-generation offspring. Analysis of double heterozygous (*Cib2^KO/+^;Whrn^wi/+^*) and double mutants (*Cib2^KO/KO^;Whrn^wi/wi^*) allowed us to determine whether deficiencies or loss of CIB2 and whirlin produces a more severe phenotype or a superimposition of pathologies.

**Figure 5:**
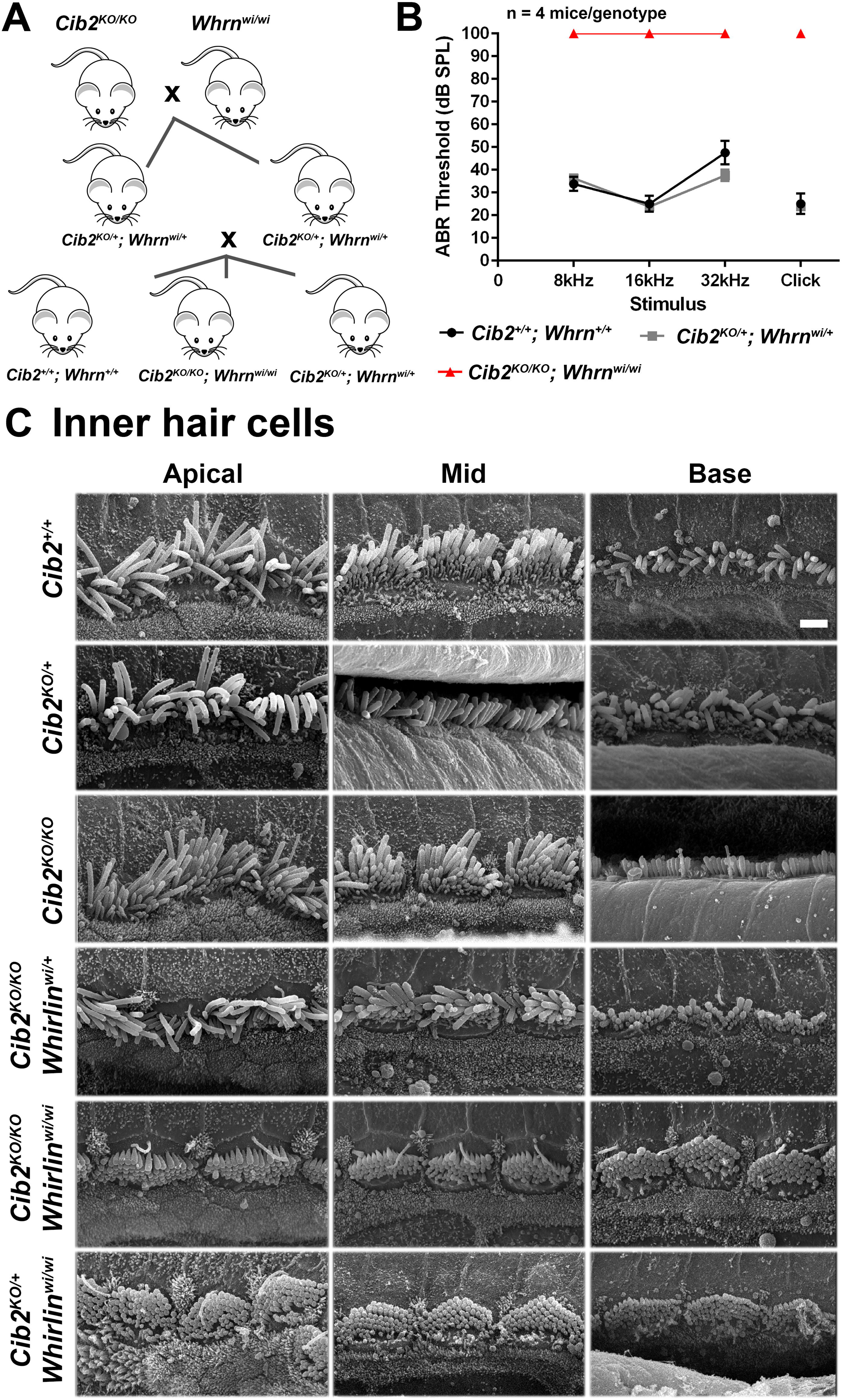
No genetic interactions between *Cib2* and *Whrn* result in hearing loss nor defects in hair cell bundle morphology. **A.** Cartoon showing the double mutant intercross breeding strategy that was employed to obtain the genotypes required for the study. Only desired genotypes are shown. **B.** Audiogram showing the ABR thresholds of 12-16 week-old mice. The data show that *Cib2^KO/+^;Whrn^wi/+^* mice (n=4) have similar thresholds to *Cib2^+/+^;Whrn^+/+^* mice (n=4), suggesting that both *Cib2* and *whirlin* are haplosufficient. As expected from ABR thresholds reported in mice homozygous for either mutation, the double homozygous mutant *Cib2^KO/KO^;Whrn^wi/wi^* mice (n=3) exhibited no response to the highest dB stimulus at any frequency tested. Data shown are mean ABR thresholds ± standard error of the mean. **C.** Scanning electron micrographs of IHCs from 2- week old *Cib2;whirler* mice. Representative scanning electron micrographs of IHC bundles from the apical, mid and basal cochlear turns of 2-week old mice. *Cib2^+/+^* and heterozygous *Cib2^KO/+^*mice have bundles that are very similar in appearance. Interestingly, in homozygous *Cib2^KO/KO^* mice IHC bundles still have kinocilia present across all turns. This developmental structure usually retracts during the first week post-partum. Moreover, additional rows of stereocilia are present compared with *Cib2^+/+^* and *Cib2^KO/+^* mice. IHC bundles of *Cib2^KO/KO^*;*whirlin^wi/+^* mice show no obvious difference from those of *Cib2^KO/KO^* mice indicating that *whirlin* haploinsufficiency does not overtly potentiate the *Cib2* null phenotype. IHC bundles of *Cib2^KO/KO^*;*whirlin^wi/wi^* mice display: short stereocilia; additional rows of stereocilia; and, kinocilia in all turns. These are features observed in both *whirlin^wi/wi^* and *Cib2^KO/KO^* mutants. IHC bundles of *Cib2^KO/+^;whirlin^wi/wi^*mice have very short stereocilia and the kinocilia is still present in the apical turn, these findings are in agreement with published findings of *whirlin^wi/wi^* where in some cases persistence of kinocilia has been noted. n ≥3 for each genotype.

**Figure 6:**
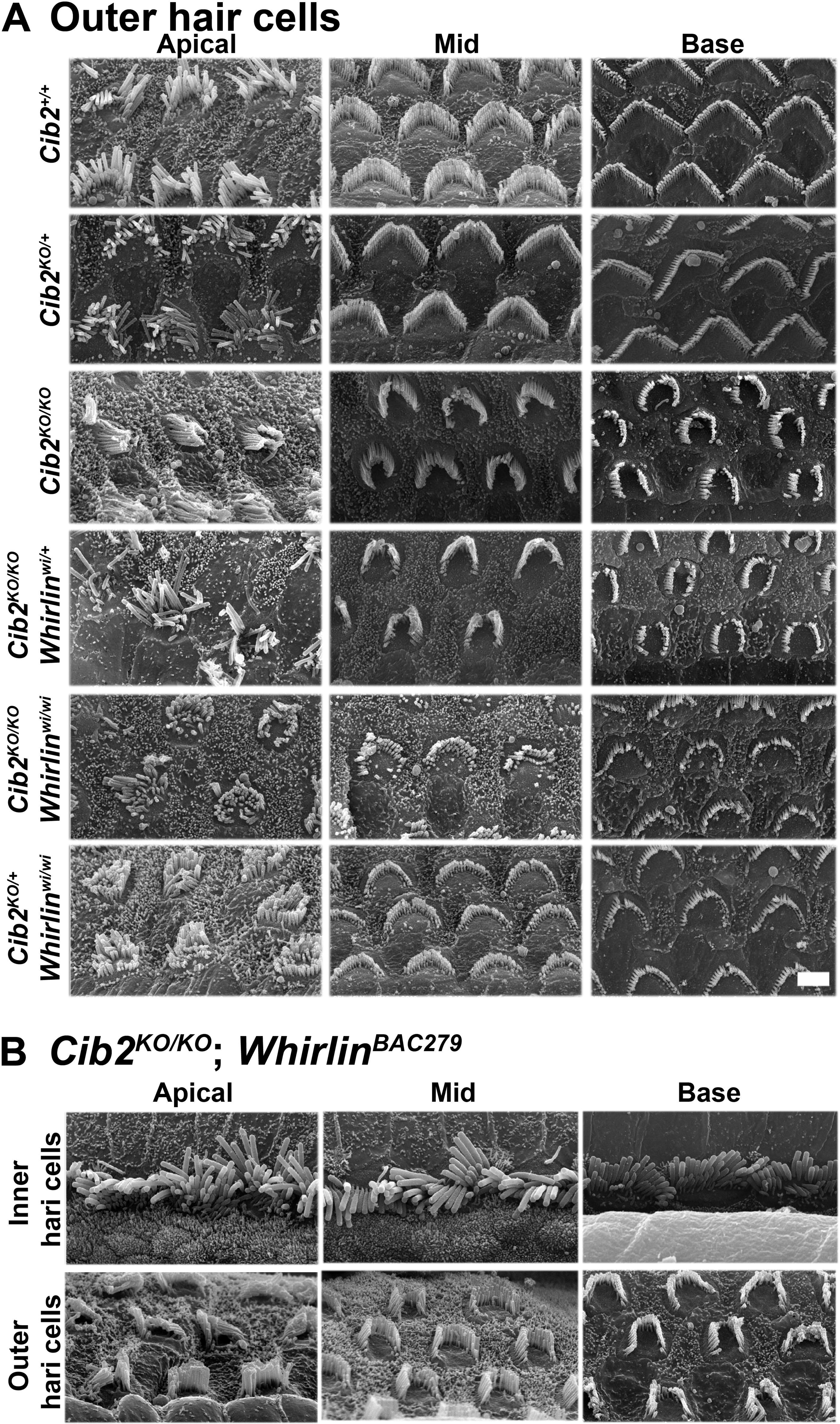
Over-expressing whirlin fails to restore stereocilia staircase pattern in *Cib2^KO/KO^* mice. **A.** Representative scanning electron micrographs of OHC bundles from the apical, mid and basal cochlear turns of 2-week old mice. *Cib2^+/+^* and heterozygous *Cib2^KO/+^* mice have bundles that are very similar in appearance. OHC bundles of *Cib2^KO/KO^* mice are poorly developed, displaying a crescent shape rather than the usual W-shape formation, and the staircase is poorly defined. Similar to IHC bundles, OHC bundles of *Cib2^KO/KO^*;*whirlin^wi/+^* mice show no obvious difference from those of *Cib2^KO/KO^*mice indicating that *whirlin* haploinsufficiency does not overtly potentiate the *Cib2* null phenotype. However, OHC bundles of *Cib2^KO/KO^;whirlin^wi/wi^* and *Cib2^KO/+^;whirlin^wi/wi^* mice are very poorly developed. n ≥3 for each genotype. **B.** Representative scanning electron micrographs of IHC and OHC bundles from the apical, mid and basal cochlear turns of 2-week old *Cib2^KO/KO^*;*whirlin^BAC279^*mice. The shape and appearance of IHC and OHC bundles appear grossly similar to those of *Cib2^KO/KO^* mice, indicating that over- expression of *whirlin* does not affect the *Cib2* null phenotype. n ≥3 for each genotype.

First wild type (WT), double heterozygous and double homozygous mutants from these crosses were subjected to auditory brainstem response (ABR) measurements at 12-16 weeks of age (Figure 5B). At this age, the double heterozygous *Cib2^KO/+^;Whrn^wi/+^*mice had ABR thresholds similar to their WT littermates (Figure 5B). In contrast to WT and double heterozygous mice, as anticipated from the reported phenotypes of *Cib2^KO/KO^* and *Whrn^wi/wi^* mice^10,10^, the double mutants *Cib2^KO/KO^;Whrn^wi/wi^*had no response to any sound stimuli (Figure 5B).

Next, we examined by scanning electron microscopy (SEM) the inner ears of double mutants and controls at 2-weeks of age. SEM images from all the double mutant mice exhibited an apparent phenotype (Figures 5B, 6A) of *Whrn^wi/wi^* mutants^17^ with some superimposition of features of *Cib2^KO/KO^* mice^10^. *Whrn* mutant had extremely short stereocilia both in inner (IHC) and outer (OHC) hair cells^17^. In contrast, the OHCs in *Cib2^KO/KO^* mutant mice had often over-growth of second row of stereocilia and horseshoe shape bundle, while the IHCs had abnormally thick third and fourth row stereocilia and persistent kinocilia^10^. The *Cib2^KO/KO^;Whrn^wi/wi^* double mutant mice had shorter stereocilia bundle with kinocilia failing to regress properly (Figures 5B, 6A). Moreover, reducing the levels of either proteins (CIB2 or whirlin)^18^ on the genetic background of other mutant strain (*Cib2^KO/KO^;Whrn^wi/+^*or *Cib2^KO/+^;Whrn^wi/wi^*) neither worsen nor rescue the normal staircase pattern in either situation (Figures 5B, 6A). Prior study has reported reduced levels of whirlin in shaft of stereocilia of auditory hair cells in *Cib2^KO/KO^* mice^19^. Therefore, we also investigated the impact of whirlin overexpression in *Cib2^KO/KO^* mice. For these studies, we crossed and generated mice that were homozygous for *Cib2^KO^*allele and were also positive for *Whrn^BAC279^* transgene^12^. Over-expressing whirlin, using the *Whrn^BAC279^* strain, also failed to restore stereocilia staircase pattern in *Cib2^KO/KO^* mice (Figure 5B). Collectively, these results support the notion that CIB2 and whirlin proteins likely have coordinated, but non- overlapping functions in orchestrating the stereocilia staircase pattern and bundle shape.

## Discussion

In this study, we demonstrate that CIB2 is multifunctional, with key independent functions in development and/or maintenance of stereocilia staircase pattern in auditory hair cells. EF2 domain of CIB2 binds to HDD2 region of whirlin and loss of CIB2 caused reduction in whirlin levels at the tips of inner hair cells stereocilia. However, double heterozygous mice from a *Cib2^KO^* x *Whrn^wi^* cross exhibited normal startle responses to sound (Figure 5B). The double homozygous mutants of *Cib2* and *Whrn* exhibited profound hearing loss. The morphology of cochlear hair cell stereocilia in double homozygous mutant mice suggest a superimposition of the phenotypes generated by each of the single homozygotes. Non-overlapping functions would be expected to generate a more pronounced phenotype. Furthermore, over-expression of whirlin in *Cib2^KO^* mice did not restore normal staircase architecture of stereocilia in cochlear hair cells. Taken together, our studies indicate that CIB2 is most likely performing a distinct function in regulating the staircase architecture of cochlear hair cell stereocilia that does not obviously overlap with the function of whirlin. The superimposition of phenotypes in the double homozygous mutant mice indicates that CIB2 and whirlin have unique and specific functions in stereocilia bundle development and patterning.

Based upon the present study, and upon the individual differences in the stereocilia bundle phenotypes of *Cib2* and *Whrn* single homozygous mutant mice^10,10^, both these genes appear to play distinct roles in establishing the correct architecture of stereocilia bundles. Recent studies have also demonstrated a critical role of MET activity in regulating the staircase pattern of stereocilia bundles in developing cochlear hair cells^20^. In Usher mutant mouse models, such as *whirler*, *shaker*, *Ush1G* and *Ush1c*, in which MET is abolished in sensory hair cells, it has been reported that the stereocilia staircase pattern is altered, and that stereocilia are dramatically reduced in length, suggesting that the MET machinery has a positive effect on F-actin polymerization^20,21^.

Several studies have reported loss of MET function in *Cib2* mutant mice^10,16,19^. However, in *Cib2* mutants the 2^nd^ and 3^rd^ row stereocilia are elongated, which is opposite to the expected <u>retraction</u> of transducing stereocilia that occurs after loss of MET current^20,21^. Thus, CIB2 likely has a role in stereocilia growth, unrelated to MET. Recent studies demonstrated that the G protein signaling modulator 2 (GPSM2) and inhibitory G proteins of the alpha family (GNAI) form a complex, which is essential for stereocilia elongation and organization into a staircase pattern^7,8^. GPSM2-GNAI binds with whirlin and the whole complex relies on MYO15A to be transported to the tips of stereocilia^7,8^. As hair cells mature, the GPSM2-GNAI complex and its partners are trafficked to the tips of stereocilia adjacent to the bare zone by the MYO15A motor, thereby establishing the “identity” of the first, tallest row of stereocilia^8^. Having in mind abnormal stereocilia heights in *Cib2* mutants, it could be speculated that CIB2 may have a role in the GPSM2-GNAI stereocilia elongation complex.

## Acknowledgements

This study was supported by NIDCD/NIH R01DC012564 (to Z.M.A) and R01DC019054 (to C.W.V.K.). The Medical Research Council MC_U142684175 (to SDMB) and MC_UP_1503/2 and MR/X004597/1 (to MRB).

## RESOURCE AVAILABILITY

### Lead Contact

Further information and requests for resources and reagents should be directed to the Lead Contact, Zubair M. Ahmed

## Materials availability

Materials generated in this study, including strains, plasmids and clones, are freely available from the Lead Contact upon request.

## Data and code availability

This study did not generate any unique datasets or code.

## Material and Methods

### Animals

All animal procedures were approved by the Institutional Animal Care and Use Committees (IACUCs) of the participating institutes. Animal strains used in this study have been previously reported^10,12^.

### Immunostaining and confocal imaging

The cochlear and vestibular sensory epithelia were isolated, fine dissected and permeabilized in 0.25% Triton X-100 for 1Ch, and blocked with 10% normal goat serum in PBS for 1Ch. The tissue samples were probed with primary antibodies against MyoXVa, whirlin, EPS8, or EPS8L1 overnight and after three washes were incubated with the secondary antibody for 45Cmin at room temperature. Rhodamine phalloidin or Alexa fluor phalloidin 488 were used at a 1:250 dilution for F-actin labeling. Nuclei were stained with DAPI (Molecular Probes). Images were acquired using either a LSM 700 laser scanning confocal microscope (Zeiss, Germany) with a 63× 1.4 NA or 100× 1.4 NA oil immersion objectives or Leica SP8 laser scanning confocal microscope with a 100x 1.44 NA objective lens. Stacks of confocal images were acquired with a Z step of 0.05-0.5Cµm and processed using ImageJ software (National Institutes of Health). Experiments were repeated at least 3 times, using at least three different animals.

### CIB2 and Whirlin constructs and plasmids

Human full length *CIB2* and WHRN cDNA constructs were generated as previously described^11^. Site directed mutagenesis was performed on the full length constructs using QuickChange PCR (Stratagene) to generate specific truncated or mutated versions. All constructs were sequence-verified before use in the experiments.

### NanoSPD assay

For NanoSPD assays, we followed the instructions reported previously^13,22^. Briefly, 60-70% confluent COS-7 cells in 6-well plates for nanoTRAP were transfected with Lipofectamine 2000 (3:1 ratio) with 1µg plasmid construct each (nanoTRAP, GFP-tagged bait, and prey), and twenty- four hours post-transfection, cells were split 1:10 ratio on glass coverslips to allow for filopodia formation. Following day, cells were fixed with 4% PFA for 15 min at RT and permeabilized with 0.2 % Triton X-100 in PBS for 15 min at RT, followed by blocking with 10% normal goat serum (NGS) in PBS for at least 30 min at RT. Primary antibodies were diluted in 3% NGS-PBS and incubated overnight at 4°C, followed by the incubations with the indicated goat secondary antibodies. A Zeiss 710 laser scanning confocal microscope or Nikon W1 spinning disk microscope was used for image acquisition.

### Co-immunoprecipitation assay

For co-immunoprecipitation (co-IP) assays, transfected HEK293T cells were washed three times with ice cold PBS and lysed in lysis buffer (150 mM NaCl, 25 mM Tris-HCl, 1% NP-40, 1 mM EDTA, 5% glycerol and proteinase inhibitor cocktails, pH 7.4) on ice for 30 min with extensively pipetting every 10 min.

The insoluble fraction was removed by centrifugation at 16,000 × g for 10 min and the lysates were split into two aliquots, one for immunoblot analysis and the other for co-IP. Equal amounts of proteins were immunoprecipitated with 25 μl GFP-Trap magnetic agarose beads (GFP-Trap®_MA, Chromotek) or anti-V5 agarose affinity gel (A7345, Sigma-Aldrich) overnight at 4 °C with gentle tumbling. The agarose beads were extensively washed four times with wash buffer (150 mM NaCl, 50 mM Tris-HCl, 0.1% NP-40, 0.5 mM EDTA, pH 7.4). The immunoprecipitated protein complexes were eluted using SDS- PAGE sample loading buffer for 5 min at 95 °C. The samples were resolved in SDS-PAGE and transferred to PVDF membranes (Bio-Rad), then subjected to western blot analysis with mouse monoclonal anti-V5-HRP antibody (1:5000, R961-25, Thermo Fisher Scientific), mouse monoclonal anti-FLAG M2-HRP antibody (1:2000, A8592, Sigma-Aldrich) or polyclonal anti-GFP-HRP antibody (1:2000, A10260, Invitrogen).

### Alphafold

An Alphafold^14^ multimer model^23^ of the CIB2/Whirlin complex was generated using the AlphaFold Colab server without template constraint (https://colab.research.google.com/github/deepmind/alphafold/blob/main/note books/AlphaFold.ipynb). Interaction interfaces were analyzed using PDBePISA (https://www.ebi.ac.uk/pdbe/pisa/)^24^. Structural models were analyzed, and figures prepared using PyMOL (The PyMOL Molecular Graphics System, Version 2.4.1 Schrödinger, LLC.)

### Auditory Brainstem Responses (ABRs)

Hearing thresholds of mice at 12-16 weeks of age (nC=C4/genotype) were evaluated by recording ABR. All ABR recordings, including broadband clicks and tone-burst stimuli at three frequencies (8, 16, and 32CkHz), were performed using an auditory-evoked potential RZ6-based auditory workstation (Tucker-Davis Technologies) with high frequency transducer RA4PA Medusa PreAmps. Maximum sound intensity tested was 100CdB SPL. TDT system III hardware and BioSigRZ software (Tucker Davis Technology) were used for stimulus presentation and response averaging.

### Scanning Electron Microscopy (SEM)

*C*ochleae were fixed in 2.5% glutaraldehyde in 0.1CM cacodylate buffer, pH 7.4 (Electron Microscopy Sciences, Hatfield, PA) supplemented with 2CmM CaCl_2_ (Sigma-Aldrich) for 1–2Ch at room temperature. Then, the sensory epithelia were dissected in distilled water, dehydrated through a graded series of ethanol, critical point dried from liquid CO_2_ (Leica EM CPD300), sputter- coated with 5 nm platinum (Q150T, Quorum Technologies, Guelph, Canada), and imaged with a field-emission scanning electron microscope (Helios Nanolab 660, FEI, Hillsboro, OR).

