## Supplemental figures for "CIB2 function is distinct from Whirlin in the development of cochlear stereocilia staircase pattern"

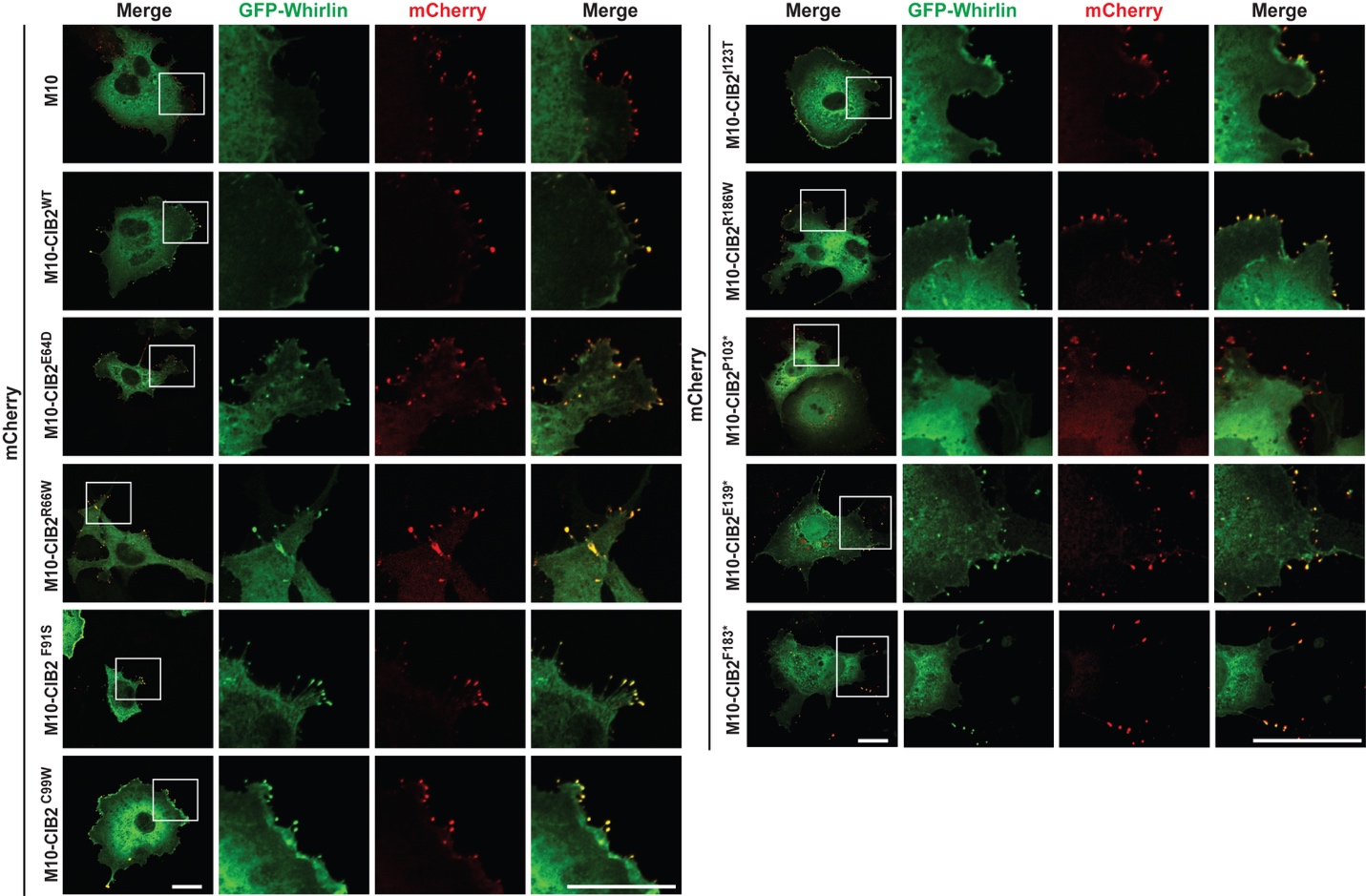

**Supplementary figure S1: Nanoscale pull down using CIB2 variants and whirlin^WT^ constructs**

COS‐7 cells were co-transfected with mCherry-myo10-CIB2 variant constructs (Baits, red) and GFP‐whirlin truncated constructs (Prey, green). Single channels are shown. mCherry-myo10 construct was used as a negative control. Scale bar = 10 μm.

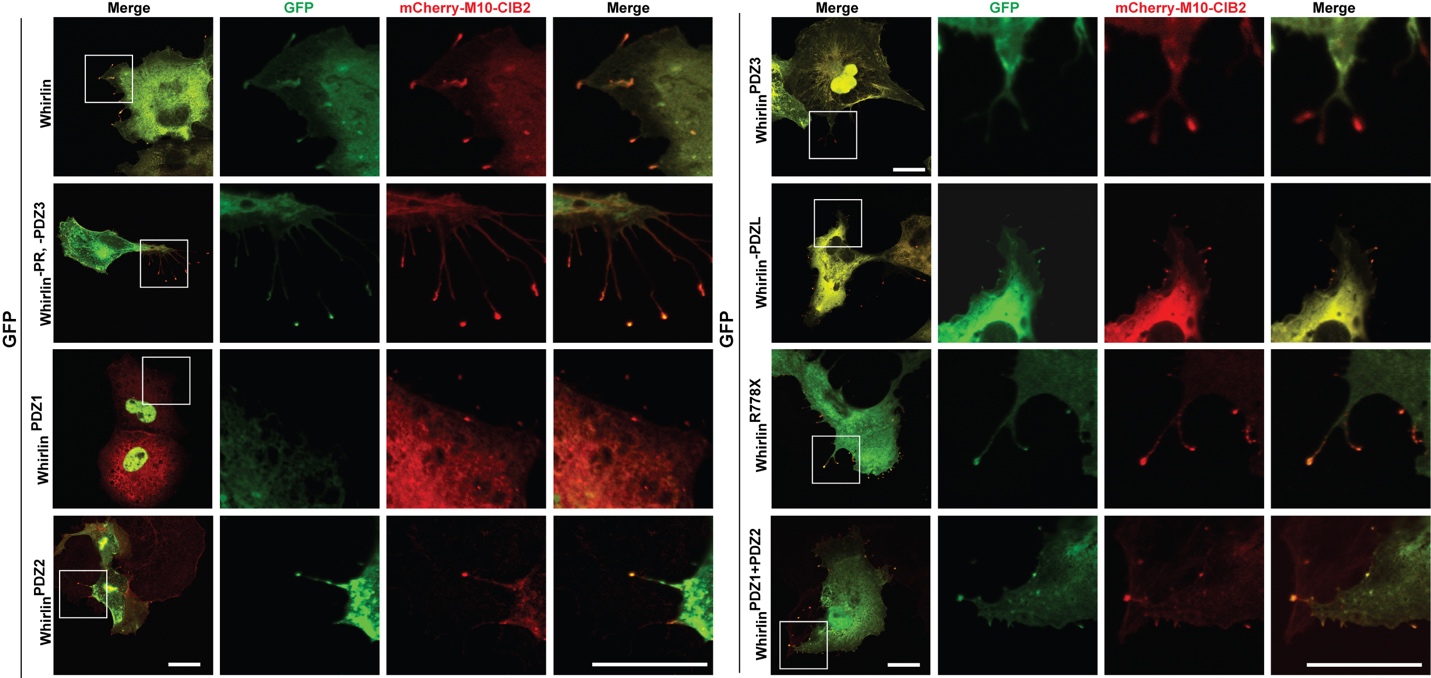

**Supplementary figure S2: Nanoscale pull down using CIB2 and truncated whirlin constructs**

COS‐7 cells were co-transfected with mCherry-myo10-CIB2^WT^ constructs (Baits, red) and GFP‐whirlin truncated constructs (Prey, green). Single channels are shown. Scale bar = 10 μm.

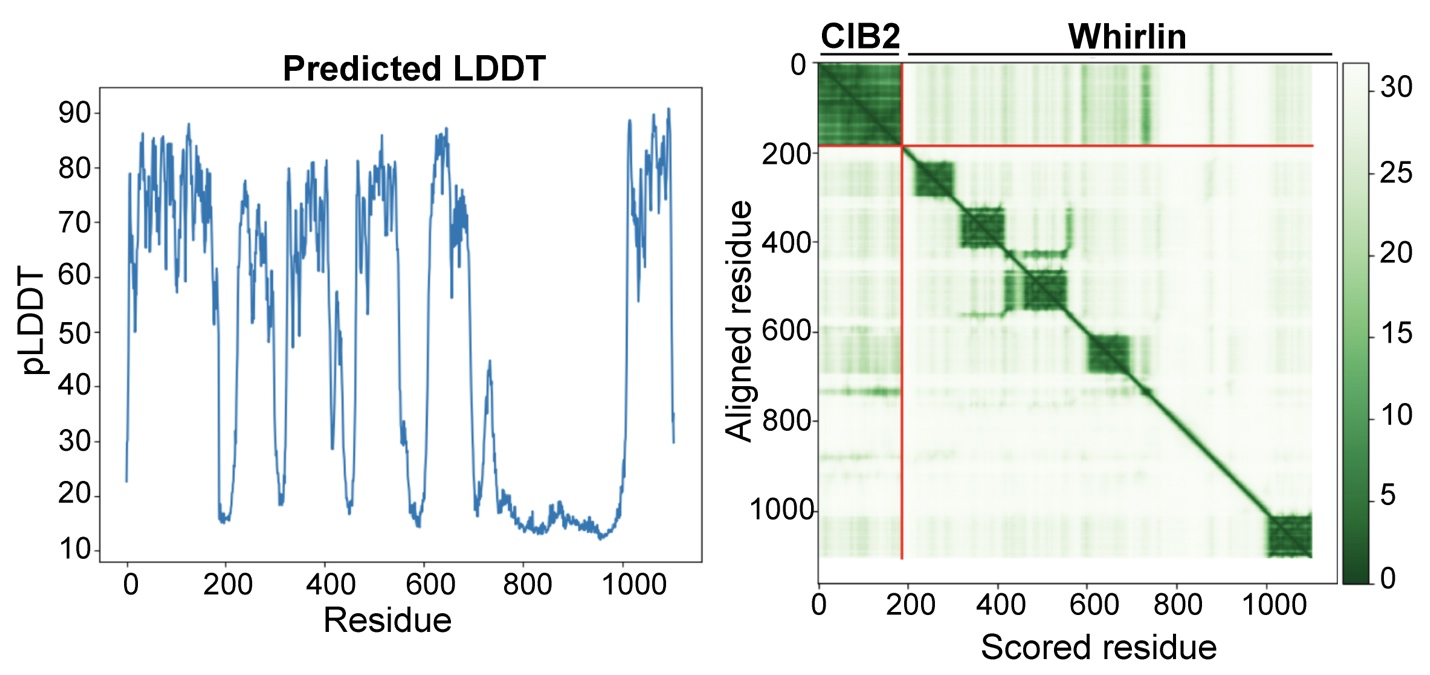

**Supplementary figure S3:** Alphafold multimer prediction quality measures for the CIB2-whirlin complex, including predicted Local Distance Difference Test (pLDDT) (left) and Predicted Aligned Error (PAE) (right).
